# Key roles of the zona pellucida and perivitelline space in promoting gamete fusion and fast block to polyspermy inferred from the choreography of spermatozoa in mice oocytes

**DOI:** 10.1101/2024.09.27.615402

**Authors:** Yaëlle Dubois, Sophie Favier, Nathan Martin-Fornier, Mohyeddine Omrane, David Stroebel, Eric Perez, Sandrine Barbaux, Ahmed Ziyyat, Nicolas Rodriguez, Christine Gourier

## Abstract

Mammalian fertilization is still lacking a comprehensive understanding of gamete fusion and block to polyspermy mechanisms. One reason is that they are highly dynamic processes involving transient events that can only be revealed and characterized by direct observation. To extend while challenging existing knowledge, this study applies real time brightfield and confocal imaging to inseminated ZP-intact mouse oocytes and statistical analyses to establish an accurate dynamic picture of the cascade of events leading to fusion and prevention of polyspermy in conditions as close to physiology as possible. These observations allow to characterize the roles of the different components of the oocyte (i.e. zona pellucida (ZP), perivitelline space (PVS), oocyte plasma membrane (OPM)) and the spermatozoon (i.e. head, flagellum) in promoting fertilization and preventing polyspermy. The kinetics we have determined challenge dogmas by showing that: (i) the first sperm is not necessarily the one that fertilizes in mice pointing to the existence of other post-penetration fertilization factors, (ii) the ZP block resulting from the cortical reaction is too slow to contribute to the prevention of polyspermy in mice. On the other hand, it evidences that the ZP directly contributes to polyspermy block in two other ways: (a) as a naturally effective entry barrier for the spermatozoon (independent of any block caused by fertilization), (b) as an effective exit barrier for components released by the OPM, which may contribute to a fast PVS block to polyspermy through the neutralizing of unwanted spermatozoa in the PVS. Moreover, our observations reveal that the ZP plays a key role in fertilization itself by channeling the flagellar oscillations of spermatozoa in the PVS to make them conducive to fusion.

## Introduction

Fertilization is the process in which haploid female and male gametes meet and fuse together to form one single cell called zygote. To fertilize, a spermatozoon must successively pass two oocyte’s barriers. The first one is the zona pellucida (ZP), the external glycoprotein envelope of the oocyte that the spermatozoon must cross to reach the perivitelline space (PVS) in which the oocyte is isolated. The second barrier is the oocyte plasma membrane (OPM) with which, once in the PVS, the spermatozoon head must interact and fuse before being engulfed by the oocyte. In mammals, only diploid zygotes resulting from the fertilization of an oocyte by a single spermatozoon can develop into a new being. When one spermatozoon successfully passes the ZP and OPM barriers, fertilization takes place. If two or more spermatozoa manage to do so, then polyspermy occurs compromising normal embryo development. To prevent such a deleterious situation, firewalls triggered by the first fertilization are deployed by the new zygote to block the other spermatozoa at one of the two oocyte’s barriers. Viable monospermic fertilization can therefore be seen as a two round game involving one oocyte and multiple spermatozoa. Round 1 consist on a rather cooperative game between the protagonists to enable fertilization. In round 2, this game turns to a race against time that the new zygote must win over all the spermatozoa that threaten its monospermic status, in order to prevent polyspermy.

Until fertilization occurs, cooperation between spermatozoa and oocyte is first reflected in the ability of spermatozoon head and oocyte’s surfaces (ZP then OPM) to adhere to each other. Adhesion to the ZP ensures that flagellum beating, which provides high motility to the spermatozoon, helps it to pass through the ZP rather than to move away from it (Baltz et al., 1988; Bleil et al., 1988; Bleil and Wassarman, 1980). Once in the narrow PVS, the spermatozoon’s flagellum movements rapidly cause its head to touch the OPM (Austin and Braden, 1956). The adhesion initiated by this contact holds the spermatozoon head to the OPM and the forces due to flagellum beating are transmitted to the oocyte. We have previously shown a correlation between a specific flagellum beating mode resulting in a pushup-like movement of its head on the OPM, and gamete fusion in contrast to other beating modes or to the mechanical inhibition of the favorable beating mode (Ravaux et al., 2016). This shows that flagellum beating is necessary, not only for locomotion, but for the fertilization event itself. However, as this was observed with ZP-free oocytes that kept the flagellum oscillation unconstrainted, the extent to which this picture is modulated in the physiological context of ZP-intact oocytes is a crucial question since the ZP imposes constraints to flagellum movements, either because the flagellum is still trapped within the ZP or because it is confined to the narrow PVS. The only chance to answer this question, directly addresses a possible role of the ZP in the fertilization event itself, is to perform real-time imaging of individual ZP-intact inseminated oocytes and to observe flagellum beating and resulting movement of the spermatozoon head on the OPM while interacting.

Once fertilization has occurred, a race against time begins between the spermatozoa, which continue on their way to fertilization, and the new zygote, which sets up firewalls to prevent them from doing so. These firewalls correspond to oocyte mechanisms, inducing changes in the zygote relative to unfertilized oocytes that result in either making it harder for spermatozoa to pass the ZP, or making the interaction between the gametes less conducive to fusion. Some post-fertilization changes observed in the ZP and OPM directly address these effects. They are referred to as ZP block and membrane block respectively. The mechanism at the origin of the ZP block is the gamete fusion-triggered exocytosis of secretory organelles (i.e. cortical granules) initially present in the cortex of unfertilized mammalian oocytes (Austin, 1956; Gulyas, 1980; Liu, 2011). This process releases proteases, glycosidases, lectins and zinc in the PVS, which act on ZP components, leading to a series of changes that are completed within half an hour to several hours, causing the ZP to lose its ability to bind to and be penetrated by spermatozoa (Baibakov et al., 2007; Barros and Yanagimachi, 1971; Bleil et al., 1981; Burkart et al., 2012; Fahrenkamp et al., 2020; Miller et al., 1993; Nishio et al., 2024; Que et al., 2017; Tokuhiro and Dean, 2018). As for the membrane block, despite the failure to identify a strong molecular basis for it in the 1990s-2000s, its existence has been clearly demonstrated by experiments in which ZP-free zygotes, challenged by a second batch of spermatozoa are shown to undergo a progressive and persistent loss of adhesion and fusion abilities that are fully established 1-2 hours after fertilization (Evans, 2020; Gardner et al., 2007; Kryzak et al., 2013; Maleszewski et al., 1996; McAvey et al., 2002; Wolf, 1978). The fact that the membrane block and the ZP block are established in approximatively the same time frame means that some (if not all) of the spermatozoa that manage to cross the ZP do so while the membrane block (and obviously de ZP block) is still not fully effective. The fact that they do not fertilize therefore suggests that firewalls are also deployed in the PVS after fertilization. This idea is supported by several observations. One of them shows that monospermic zygotes with motile spermatozoa in their PVS, with which they are apparently unable to fuse, can nevertheless be successfully re-fertilized after removal of their ZP (and thus their removal from the PVS) if they are re-inseminated less than one hour after the first fertilization (Maluchnik and Borsuk, 1994). Another study reports that a spermatozoon in the PVS of a zygote cannot fertilize more than 1 min after initial fertilization (Sato, 1979). For a brief period in the 1990s, the PVS block was hypothesized to be due to some cortical granule contents that remain in the PVS after cortical granule exocytosis, forming the so-called cortical granule envelope. Like the ZP, it was thought to act as a physical barrier for spermatozoa reaching the PVS (Dandekar and Talbot, 1992; Talbot and Dandekar, 2003). However, further study has ruled out its involvement in the PVS block, marking, the end of research on this subject (Hoodbhoy et al., 2001). Today, we know that the zygote releases oocyte membrane proteins JUNO and CD9 in the PVS shortly after fertilization (Bianchi et al., 2014; Ravaux et al., 2018). Depleting the OPM from the two oocyte membrane proteins identified as essential for gamete fusion in mice while releasing them in the PVS, provides food for thought to re-launch research into both the membrane block and the PVS block (Bianchi et al., 2014; Kaji et al., 2000; Le Naour et al., 2000; Miyado et al., 2000; Ravaux et al., 2018).

Fertilization and block to polyspermy are highly dynamic processes. They involve transient events in which multiple components of the spermatozoon and oocyte interact simultaneously. The only way to reveal and characterize these events is to observe them directly under conditions as close to physiology as possible. To this end, this study applies real-time brightfield and confocal imaging to dozens of individually imaged inseminated ZP-intact mouse oocytes. From the moment a spermatozoon begins to cross the ZP of an oocyte, its movements and progression reflect the interactions of its head and flagellum with the different compartments of the oocyte (ZP, PVS, and OPM). The visual information we obtained from more than a hundred spermatozoa, correlated with a statistical analysis of the time of their penetration into the PVS and, when relevant, the time of their fusion, allow us to establish a precise dynamic picture of the sequence of events leading to fusion and prevention of polyspermy, and of the role of each gamete component in promoting fertilization and preventing polyspermy. The kinetics we have determined challenge dogmas on the role of the ZP in blocking polyspermy in mice, and on the omnipotence of the first spermatozoon to penetrate the oocyte. The new information they provide on the actual roles of the ZP and of the PVS lead us to shake some classical views, opening up new avenues of research.

## Results

Throughout this study, we will discuss two different events involved in fertilization that are often referred indifferently in literature as “penetration”. The first event is the entry of a spermatozoon into the PVS of an oocyte after having crossed its zona pellucida (ZP). The second event refers to the fusion of the spermatozoon head with the oocyte plasma membrane (OPM) followed by its internalization into the oocyte cytoplasm. To avoid any confusion, we decided to reserve the term “penetrate” to the first event and the term “fertilize” to the second one. Accordingly, a spermatozoon crossing the ZP and entering the perivitellin space (PVS) is designated as a “penetrating spermatozoon”. If it additionally fuses with the OPM, it also becomes a “fertilizing spermatozoon”.

### Multi-penetration and polyspermy under *in vivo* conditions and standard and kinetics *in vitro* fertilization conditions

The amount and timing of the spermatozoa reaching an oocyte are parameters that likely influence the number of penetrations, of fertilizations and thus the capacity of a zygote to develop normally. These parameters can vary a lot from *in vivo* to *in vitro* insemination conditions, or between different *in vitro* inseminations conditions. In order to determine the parameters controlling penetrations, fertilizations, and prevention of polyspermy, we conducted a comparative study on more than 500 oocytes inseminated in 3 different conditions. In condition 1, 211 oocytes were recovered from females 15 hours after mating and were therefore submitted to the so-called “physiological *in vivo”* insemination conditions. In condition 2, 220 oocytes were submitted to the so-called “standard *in vitro”* fertilization conditions in which cumulus/oocytes masses were inseminated with 10^6^ capacitated spermatozoa/mL for 4 hours. In condition 3, 93 oocytes were submitted to the so-called “kinetics *in vitro”* experiments in which the oocytes with the spermatozoa bound to their ZP were removed from standard *in vitro* fertilization conditions 15 minutes after insemination to be filmed in close-up during the next 4 hours.

*In vivo* fertilization experiments provide reference values regarding penetration and fertilization of oocytes in physiological conditions, where the ejaculated spermatozoa reach the ovulated oocytes rather sparingly over time (Figure 1)(Austin and Braden, 1952). They reveal that despite such a natural regulation, multi-penetration is frequent among the fertilized oocytes: 16 ± 6% are penetrated by 2 to 3 spermatozoa (Figure 1A-C). Yet, all of them are monospermic (Figure 1B). This, indicates the existence of an efficient post-penetration contribution to the polyspermy block ensuring that only one of the penetrated spermatozoa fertilizes. However, such a post-penetration contribution to the polyspermy block is physiologically useful only if monospermic multi-penetrated oocytes are able to develop normally, which to our knowledge has never been proven in mice. We set out to verify this by implanting in 1 pseudo-pregnant female (female mated with a sterile male), 16 two-cells stage monospermic zygotes with additional unfertilizing-spermatozoa in their PVS and in 2 pseudo-pregnant females, 16 two-cells stage monospermic zygotes per female with no extra unfertilizing spermatozoon. The absence of differences in litter sizes (6, 6 and 7 pups) confirms that the presence of spermatozoa in the perivitelline space of monospermic oocytes is not an obstacle to normal embryo development, while underlining the physiological importance of the fertilizing block process preventing polyspermy in multi-penetrated oocytes.

**Figure 1:**
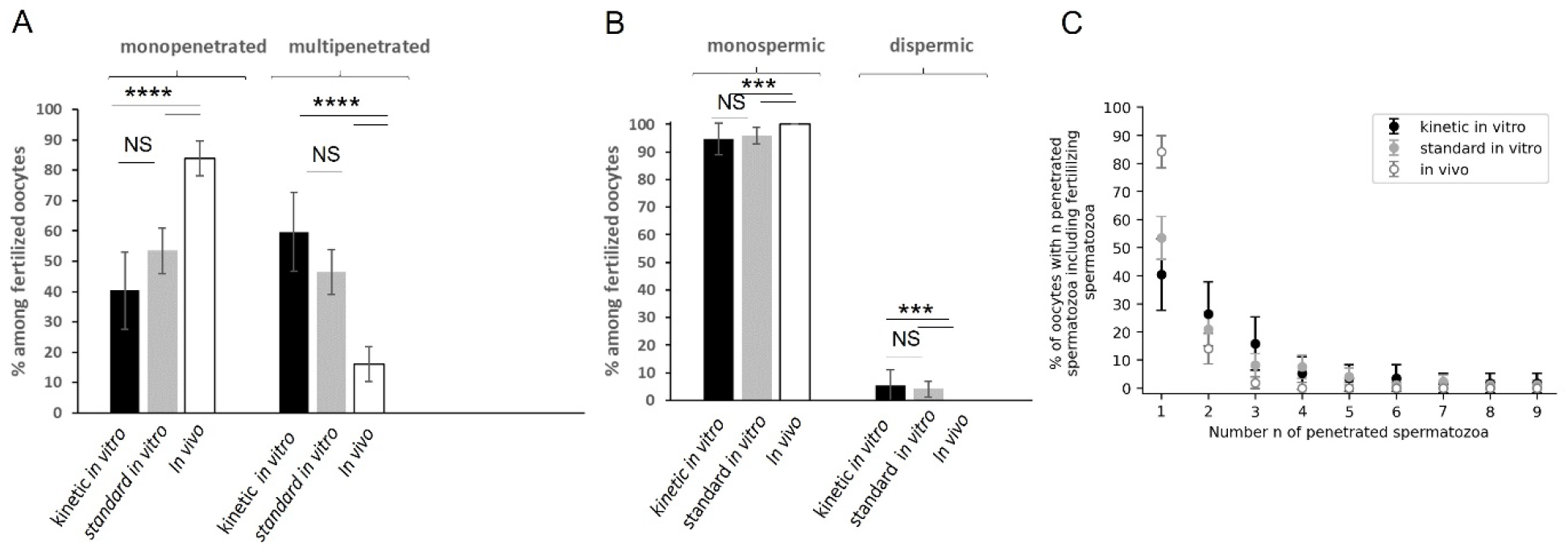
Multi-penetration and polyspermy during *in vivo* fertilization and standard and kinetic *in vitro* fertilization. A –rates of mono- and multi-penetrated fertilized oocytes B-rates of monospermic and polyspermic oocytes C-rate of fertilized oocytes with n penetrated spermatozoa. Error bars corresponds to 95% confidence interval.

Standard *in vitro* fertilization conditions differ from in vivo fertilization conditions in that oocytes are rapidly surrounded by numerous spermatozoa that can simultaneously bind to their ZP and be continuously relayed by others if they detach. In this situation of excess spermatozoa, the main limit to the number of penetrations comes from the ZP itself and the temporal evolution of its permeability to spermatozoa. We found that multi-penetration affects 47 ± 8% fertilized oocytes (Figure 1A), with the number of spermatozoa penetrating these oocytes ranging from 2 to 9 (Figure 1C). These figures are low compared to the huge number of spermatozoa available for penetration, showing that ZP effectively limits penetration. However, as far as the risk of polyspermy is concerned, these figures remain very high, since beyond one penetrated spermatozoon, prevention of polyspermy depends solely on the inability of all but one to interact properly with the OPM to end in gamete fusion. Interestingly, we observed that 96 ± 3% of oocytes remain monospermic, and the few cases of polyspermy are limited to dispermy only (Figure 1B), which confirms *in vitro* the existence of a fertilization block preventing polyspermy downstream of the ZP.

Kinetic *in vitro* fertilization conditions differ from standard *in vitro* conditions in that the number of spermatozoa likely to penetrate is limited to the spermatozoa already bound to the ZP of the oocyte when they were isolated (i.e. typically a few dozen) 15 minutes after insemination. In terms of cumulated number of spermatozoa reaching an oocyte over time, kinetic *in vitro* conditions are therefore probably closer to physiological *in vivo* fertilization conditions than to standard *in vitro* conditions where the reservoir of spermatozoa likely to enter in contact with the oocyte is almost unlimited. However, in terms of the number of spermatozoa rapidly and simultaneously bound to the ZP of an oocyte, kinetic *in vitro* conditions are identical to standard *in vitro* ones and very different from *in vivo* conditions where sperm reach the oocytes sparsely over time. The fact that (i) the rates of mono- and multi-penetrated fertilized oocytes (Figure 1A), (ii) the distribution of the number of spermatozoa in these oocytes (Figure 1C), and (iii) the rates of mono-and di-spermic oocytes (Figure 1B) are similar for standard and kinetic *in vitro* fertilizations, but significantly different from *in vivo* fertilizations, suggests that multi-penetration and polyspermy depend more on the number of spermatozoa rapidly and simultaneously bound to an oocyte than on the cumulative number of spermatozoa reaching an oocyte over time in agreement with what will be discussed further below.

### Kinetics of sperm engulfment after fertilization and release of the second polar body

Gamete fusion is the trigger of various processes affecting the oocyte, including engulfment of the fertilizing spermatozoon and release of the second polar body (PB2) from the oocyte, marking the end of meiosis. Unlike *in vivo* and standard *in vitro* fertilizations which are performed blindly, the experimental conditions of kinetic *in vitro* fertilization provide access to temporal and visual information about the main fertilization related events. Eight fertilizations for which we were able to monitor in real time the sudden and definitive arrest of the flagellum movements -a sign of imminent gamete fusion (Ravaux et al., 2016)-(Video 1 and Video 3, white star fertilization event flag), allowed us to establish an average post-fertilization kinetics of the spermatozoon engulfment and PB2 release (Figure 2).

**Figure 2:**
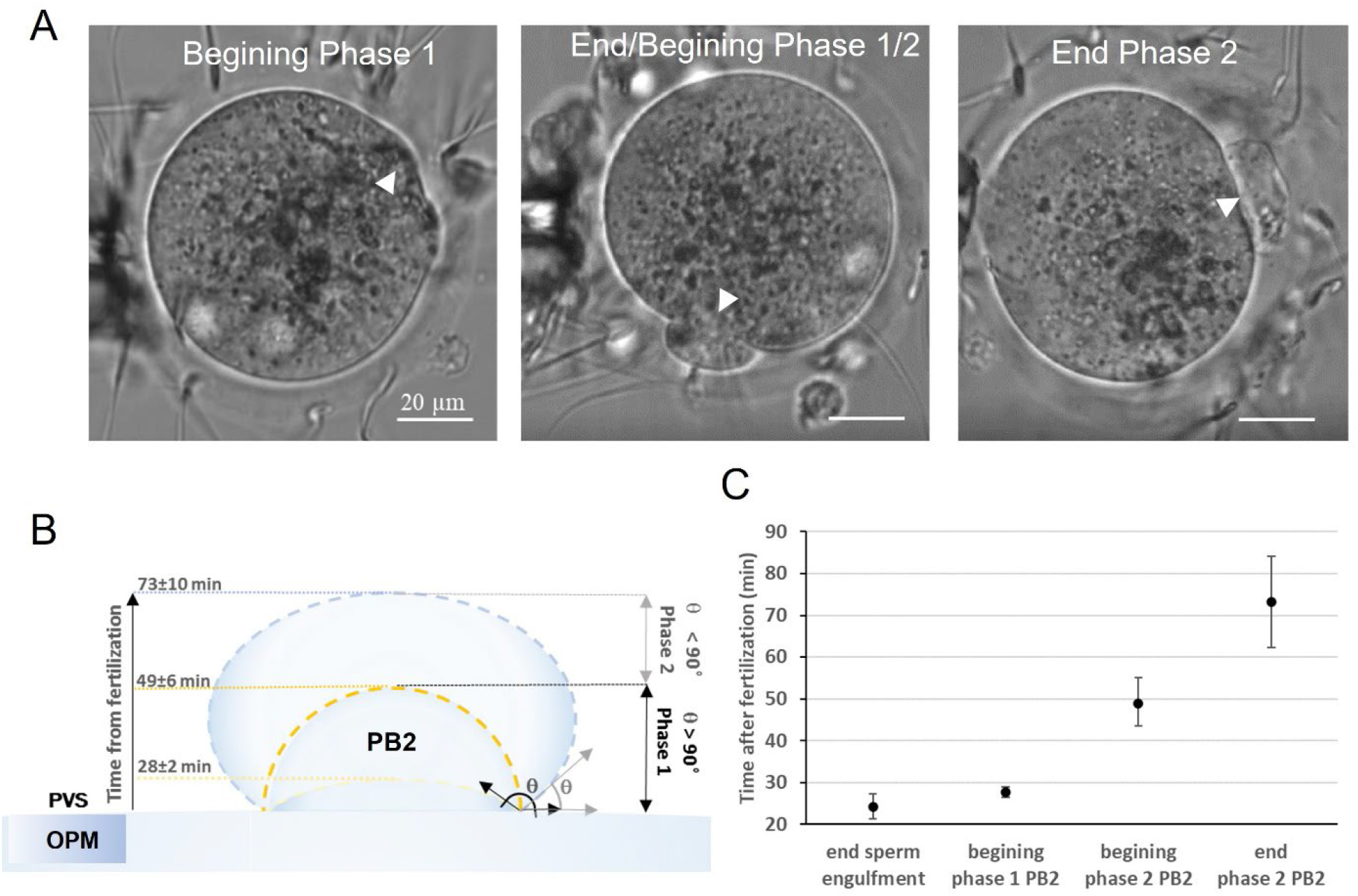
Kinetics of post-fertilization sperm engulfment and second polar body release. A-sequences from the beginning of Phase 1 to the end of Phase 2 (using the geometric criteria described in B for distinguishing phase 1 and 2) of PB2 release. White arrow heads indicate the position of the second polar body. B-geometric criteria used to distinguish between Phase 1 and 2 of PB2 release: Phase 1 as long as the angle between the OPM and PB2 is higher than 90° and stage 2 when this angle starts to become smaller than 90°. C-mean time ± standard deviation of sperm engulfment and PB2 release after fertilization.

Through bright field observations, the engulfment of the fused spermatozoon is reflected by a progressive decrease in the visibility of its head on the membrane, to the point where, completely engulfed, it can no longer be distinguished at all, which happened typically 24±3 min (mean ± standard deviation) after fusion (Figure 2C). Then the release of PB2 starts, reflected by the formation of an increasing rounding hernia on the OPM surface (Figure 2A). In this long release process, we chose to distinguish two phases: phase 1 as long as the angle between the OPM and PB2 is higher than 90° and phase 2 when this angle starts to become smaller than 90° (Figure 2B). With these criteria, phase 1 of PB2 is found to begin 28± 2min (mean ± SD) after gamete fusion and to end with the beginning of phase 2, 49± 6 min after gamete fusion, while phase 2 ends when PB2 release is completed 73 ±10 min after fusion (Figure 2C).

### Kinetics of penetrations and fertilizations

Over the 93 oocytes which were subject to kinetic *in vitro* experiments, 57 were penetrated by a total of 138 spermatozoa and fertilized by 60 of them, 14 were penetrated by 46 spermatozoa but not fertilized, and 22 were not penetrated at all. For all these spermatozoa, we experimentally determined a penetration time chosen as the time at which the head emerged from the ZP in the PVS as well as a fusion time (when applicable) corresponding to the time at which the penetrated spermatozoon in interaction with the OPM suddenly and definitely stopped beating. These data are reported in two chronograms, one for the fertilized oocytes (Figure 3A) and the other for the penetrated but not fertilized oocytes (Figure 3B). The inaccuracy on the determination of a penetration time or fertilization time (symbolized by the length of the time windows in figure 3) is inherent to our experimental protocol where oocytes are filmed in turn. The accuracy is high (very short time window in Figure 3A-B) when penetration or fertilization occurs during one round of observation of the oocyte and is therefore seen live. It decreases (larger time window in Figure 3 A-B) when occurring between two rounds of observation of the oocyte. In this latter case, the best accuracy we can get on the penetration time is the gap between the end of the round preceding penetration and the start of the following round when the presence of a new spermatozoon in the PVS is noted. As for a fertilization event occurring between two rounds of observations, the accuracy on its fertilization time is not necessarily as coarse as the gap between these two rounds since fertilization is followed by the engulfment of the fertilizing spermatozoon and the release of PB2 whose stages can easily be evaluated during the successive rounds of observations of the oocyte and correlated to the previously established post-fertilization timing, providing relevant landmarks to bound the fertilization time window.

**Figure 3:**
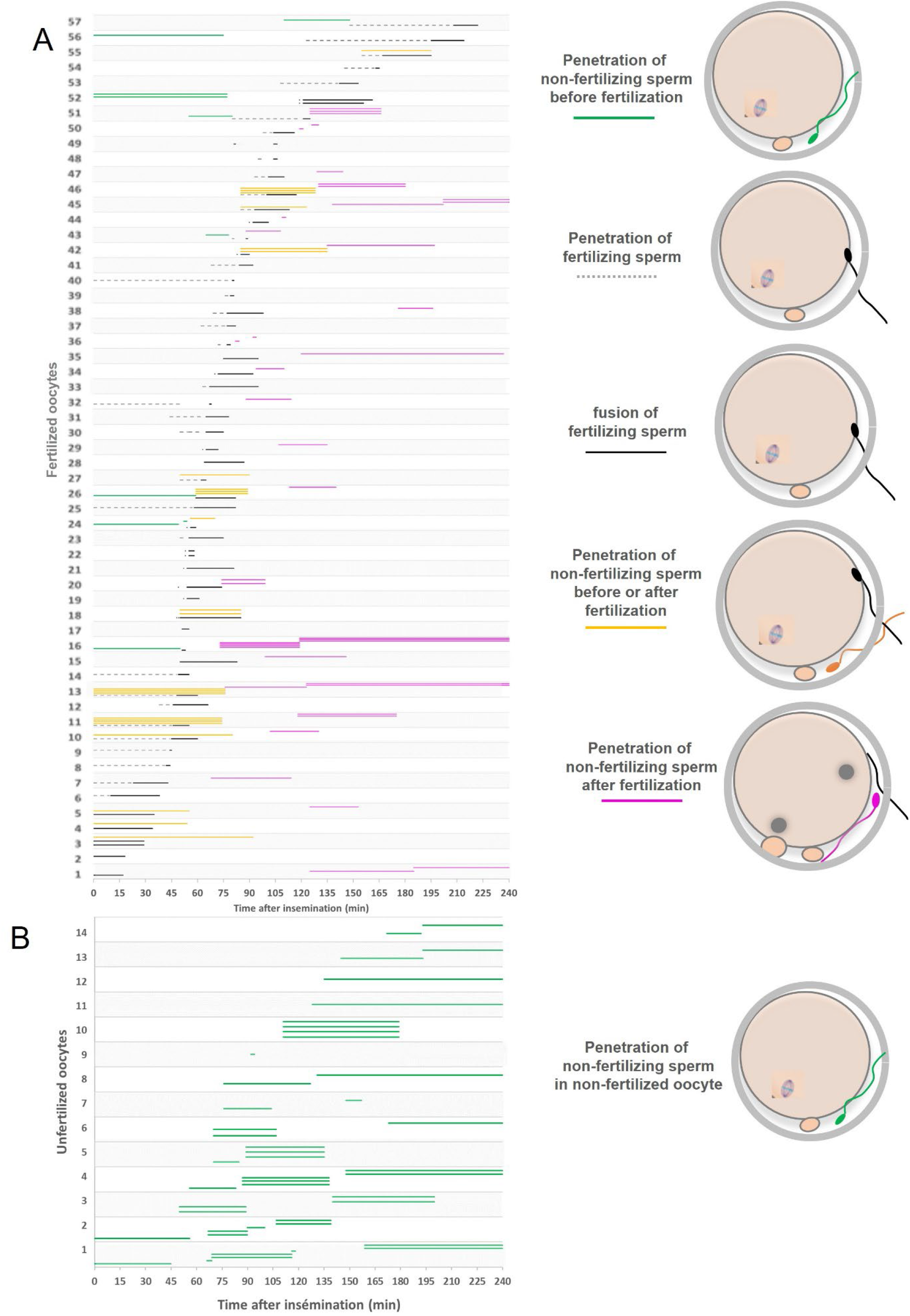
Post-insemination timelines of penetrations and fertilizations. A-Fertilized oocytes (57 oocytes, 138 penetrated spermatozoa including 60 fertilizing spermatozoa). Each strip corresponds to one oocyte. Each dashed black line within one strip corresponds to the post-insemination time window during which a fertilizing spermatozoon penetrated the oocyte, and the full black line within the same strip corresponds to the time window during which this spermatozoon fused with the OPM. Each green, yellow and purple lines within the one strip represents the penetration time window of an additional non-fertilizing spermatozoon. Green and purple colors are visual helps to indicate that penetration of the non-fertilizing spermatozoon occurred before (green) or after (purple) the penetration and fusion of the fertilizing spermatozoon. Yellow color means that we can neither determine which of the unfertilizing or fertilizing spermatozoon penetrated first, nor which of penetration of the unfertilizing spermatozoon or fusion of the fertilizing spermatozoon occurred first since the penetrations and the fertilization of these spermatozoa all took place during the same blind gap between two rounds of observation of the oocyte. B-Unfertilized penetrated oocytes (14 oocytes, 46 penetrated spermatozoa). Each strip corresponds to one oocyte. Each green line within one strip represents the post-insemination time window of an unfertilizing spermatozoon.

### The first penetrated spermatozoon is not necessarily the fertilizing one

The penetration and fertilization chronogram of fertilized oocytes show that the first penetrated spermatozoon does not necessarily fertilize (Figure 3A). This result contradicts an old dogma based on the assumption that any spermatozoon able to reach an oocyte and cross the ZP (and a fortiori the first of them) has the intrinsic skills to fertilize. However, confronted to the systematic determination of the order of penetration and fertilization of spermatozoa, this dogma falls apart. Here we find that all the green spermatozoa (10 spermatozoa) (Figures 3A) penetrated their oocyte (8 oocytes) before the fertilizing spermatozoa (60 fertilizing spermatozoa in 57 oocytes) and so did an unknown number from 0 to 26 of yellow spermatozoa. In the end, we determined that 61,7% of the fertilizing spermatozoa were the first to penetrate their oocyte, 18.3% were not and uncertainty remains over the last 20% of fertilizing spermatozoa (Figure 4A).

**Figure 4:**
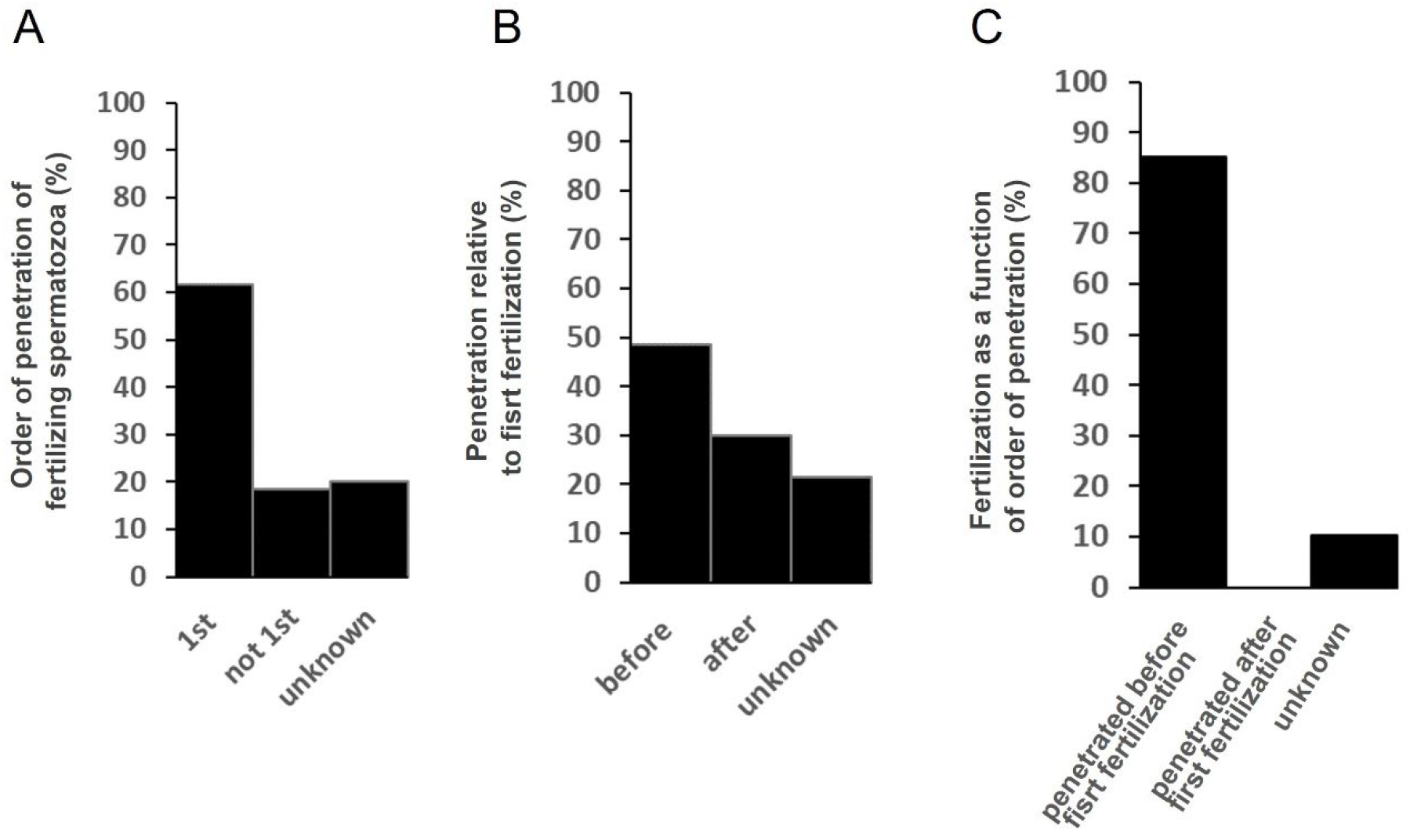
Fertilization and order of penetration in fertilized oocytes. A-order of penetration of fertilizing spermatozoa. B-rates of spermatozoa penetrated before and after first fertilization. C-rates of fertilizing spermatozoa as a function of their order of penetration.

### Spermatozoa can penetrate after the first fertilization but have very low chances to fertilize

Of the 138 spermatozoa that penetrated the 57 fertilized oocytes, 48.6% penetrated before first fertilization and 85.1% of them fertilized, 30% penetrated after first fertilization and none of these fertilized, and of the last 21.4% penetrating spermatozoa -for which there is uncertainty about the time of their penetration relative to the time of first fertilization-10.3% fertilized (Figure 4B-C). These results show that a spermatozoon that penetrates a yet unfertilized oocyte has high chance of fertilizing it, but its chance drops sharply if another spermatozoon (whether penetrated before or after it) manages to fertilize it first. As for the many spermatozoa penetrating after the first fertilization, their chance to fertilize is almost zero. This suggests that first fertilization contributes effectively to the fertilization block, but less so to the penetration block.

### Statistical treatment of penetrations and fertilizations chronograms to study penetration and fertilization blocks

To take advantage of our large sample size and make our quantitative analyses more informative and precise, we have implemented a statistical analysis to process the penetration and fertilization time windows obtained experimentally. This statistical analysis is based on the assumption that the probability distribution of penetration or fertilization is uniform within a given time window and takes into account for all oocytes all the possibilities of penetration and fertilization (when relevant) times of one spermatozoon in an oocyte relatively to all the other spermatozoa from the same oocyte. In the following, all values derived from this statistical analysis are given with a 95% confidence interval. The subscripts *PP*_1_, *PF*_1_ *NF, PP* and *PolyB* refer to post-first penetration, post-first fertilization, no fertilization, penetration block and polyspermy block respectively.

### The penetration block

To characterize the kinetics with which penetration of spermatozoa in the PVS falls down after a first fertilization, we determined the post-first fertilization penetration index 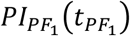 (i.e. mean number of spermatozoa per oocyte that have penetrated an oocyte between the first fertilization time (time zero) and time 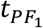 and its uncertainty 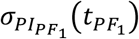 (Figure 5A). 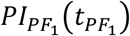 tells us how, in average, the number of penetrating spermatozoa increases over time in an oocyte after an initial fertilization. 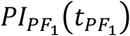 is fitted by the function:

**Figure 5:**
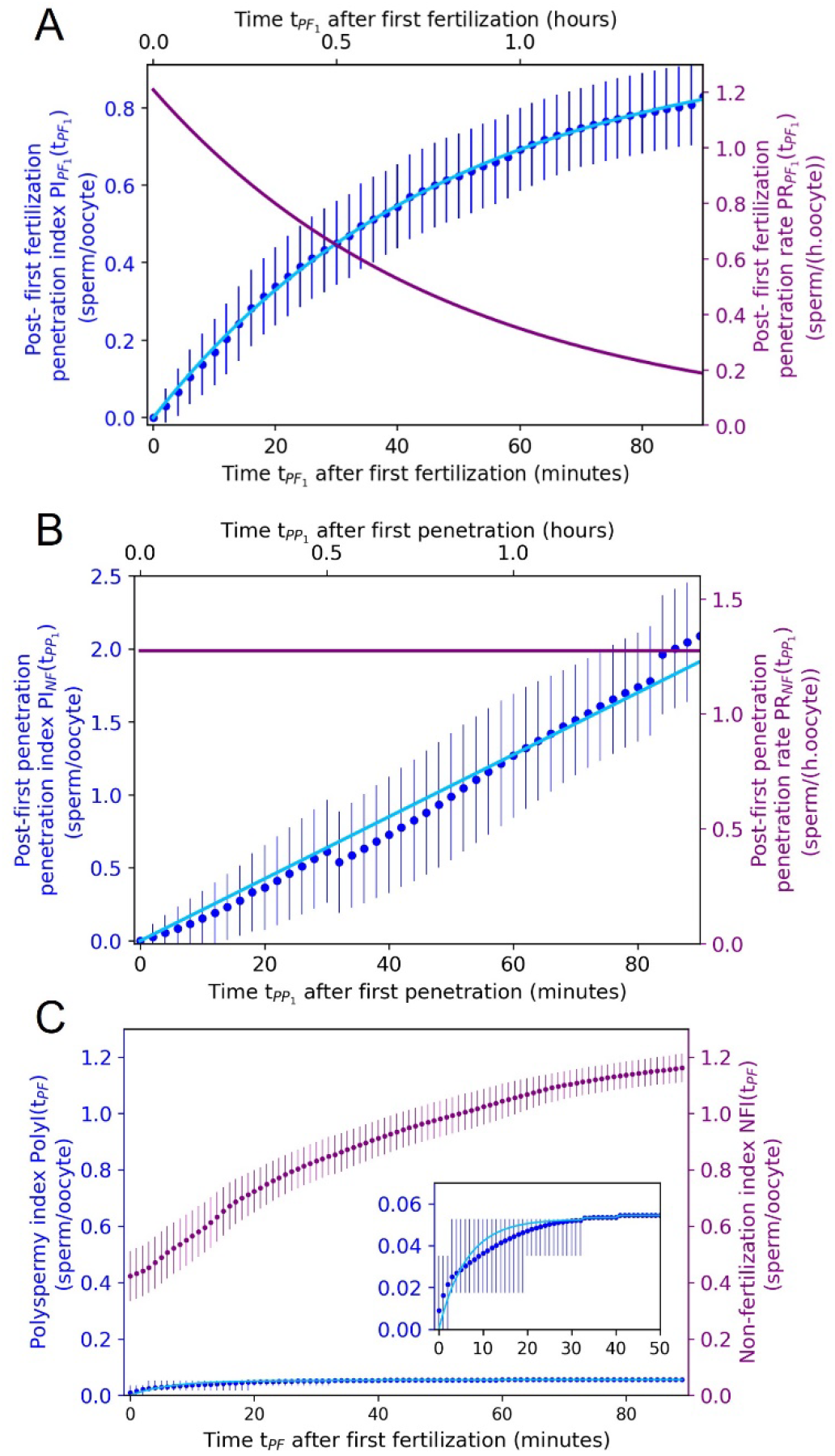
Kinetics of penetration and polyspermy blocks. A-Post-first fertilization penetration index and its uncertainty (dark blue dots and error bars) obtained by statistical analysis of fertilized oocytes’ penetration and fertilization chronogram (Figure 3A). Its fitting curve (light blue line) shows that after the first fertilization the ZP becomes impermeable to spermatozoa with a time constant τ_PB_ = 48.3 ± 9.7 min and in average, 0.97 ± 0.10 sperm/oocyte can still penetrate. Post-fertilization penetration rate (purple line) obtained by derivation of the post-fertilization penetration index fitting function gives the mean post-fertilization time evolution of the permeability of the ZP to spermatozoa. Starting from a penetration rate equal to 1.21 ± 0.37 sperm/oocyte/hour corresponding to the “natural” permeability of ZP to spermatozoa, it decreases after a first fertilization up to become impenetrable with a block to penetration time constant τ_PB_ = 48.3 ± 9.7 min. B-Post-first penetration penetration index and its uncertainty (dark blue dots and error bars) obtained by statistical analysis of unfertilized oocytes’ penetration chronogram (Figure 3B). It is fitted by a linear function (light blue line) showing a steady penetration rate equal to 1.28 ± 0,07 sperm/oocyte/hour (purple line). The steady penetration rate shows that penetration has no influence on ZP permeability to spermatozoa. C Polyspermy index and its uncertainty (dark blue dots and error bars). Insert is a zoom of the 50 first minutes of the polyspermy index. Its fitting curve (light blue line) shows that after first fertilization, the polyspermy block takes place with a time constant τ_PB_ = 6.2 ± 1.3 min. Polyspermy index and non-fertilization index curves were obtained by statistical analysis of fertilized oocytes’ penetration and fertilization chronogram (Figure 3A).

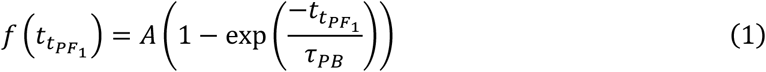

It reaches a plateau *A* = 0.97 ± 0.10 spermatozoon/oocyte with a time constant *τ*_*PB*_ = 48.3 ± 9.7 minutes. After the first fertilization, the penetration rate 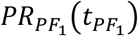, given by the slope of 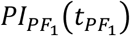, slows down to zero with the time constant *τ*_*PB*_, from an initial value at 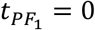 (before the onset of any fertilization-related contribution to the penetration block) equal to 1.21 ± 0.37 sperm/hour/oocyte. As we have shown that the number of available spermatozoa for penetration is not a limiting factor under our *in vitro* conditions, the decrease measured in penetration rate after first fertilization must result from a progressive decrease in the ZP permeability to spermatozoa occurring with a time constant *τ*_*PB*_ = 48.3 ± 9.7 minutes. We can assume that the trigger for this reduction is the first fertilization of the oocyte, but at this stage we cannot rule out the possibility that the permeability of the ZP began to decrease before fertilization, notably because of penetrations (including the penetration of the first fertilizing spermatozoon), or binding of spermatozoa to the ZP or any other possible event that may impact the ZP before fertilization.

To answer this question, we took advantage of the 14 oocytes that were multi-penetrated but not fertilized (Figure 3B). Although their fertilization failure highlights a fertilization defect, spermatozoa can penetrate these oocytes and if penetration itself triggers any evolution in ZP permeability, it should be reflected in the temporal evolution of the mean penetration rate derived from the mean penetration index of these oocytes. Taking for each oocyte the first penetration time as the origin of time 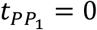, we have therefore applied our statistical analysis to the chronogram of these oocytes to obtain the evolution of the post-first penetration penetration index (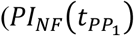 (i.e. mean number of spermatozoa per oocyte that penetrated between the first penetrating spermatozoon (time 0) and time *t*_*PP*_) and its uncertainty (Figure 5B). 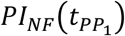 is fitted by the linear function

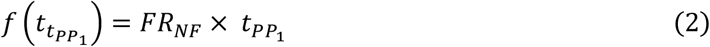

The best fit is obtained for a steady penetration rate *FR*_*NF*_ equal to 1.28 ± 0,07 sperm/oocyte/hour (Figure 5B). This steady penetration rate shows that neither penetrations nor other events taking place before fertilization impact ZP permeability. It also proves in retrospect that the post-fertilization decrease in ZP permeability observed on fertilized oocytes was indeed the result of fertilization alone. Finally, the similar values of the steady penetration rate of unfertilized oocytes (1.28 ± 0,07 sperm/oocyte/hour) and of the penetration rate at time of first fertilization of fertilized oocytes (1.21 ± 0.37 sperm/oocyte/hour at 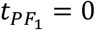) confirm a normal ZP permeability of the unfertilized oocytes, while providing a quantification of an important characteristic of the ZP: its original permeability to spermatozoa. Its low value makes the ZP an effective barrier to penetration even before fertilization takes place, limiting the risk that several spermatozoa penetrate simultaneously and therefore the risk of polyspermy.

Overall, these data reveal that penetration block is due first to the low natural permeability of the ZP to spermatozoa (∼1.3 sperm/oocyte/hour in average), to which is added, from the first fertilization time, a progressive and persistent loss of ZP permeability, triggered by fertilization itself and referred in literature as the ZP-block, taking place with the time constant *τ*_*PB*_ = 48.3 ± 9.7 minutes. Before the ZP becomes fully impermeable, in average *AA* = 0.97 ± 0.10 spermatozoon per oocyte can still penetrate.

### The polyspermy block

To characterize the kinetics with which fertilization by new penetrating or already penetrated spermatozoa falls down after a first fertilization (i.e. polyspermy block), we determined the polyspermy index 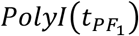 (i.e. mean number of polyspermic fertilizations per oocyte at time 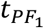 after the first fertilization of the oocyte) and its uncertainty 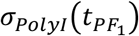 from the statistical treatment of the fertilization time windows in the 3 dispermic oocytes (Figure 5C). 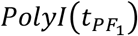 is fitted with the function

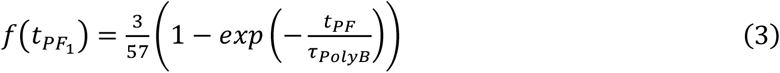

being the number of polyspermic oocytes, and 57 the number of fertilized oocytes) and the best fit is obtained with *τ*_*PolyB*_ = 6.2 ± 1.3 minutes. Following computation of the penetrated non-fertilizing spermatozoa at time 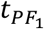, we determined the non-fertilization index 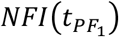 (i.e. mean number of available spermatozoa in the PVS at time 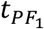 after the first fertilization of the oocyte) which shows that in addition to the 3 spermatozoa that actually fertilize after at first fertilization, numerous penetrated spermatozoa are available in the PVS of the oocytes but do not fuse. *τ*_*PolyB*_ = 6.2 ± 1.3 minutes can thus be interpreted as the time constant of the polyspermy block.

#### Fertilization causes de-adhesion of spermatozoa from the ZP

A rough counting of the number of spermatozoa linked to the oocyte ZP was carried out during the oocyte observation rounds. Level 4 is attributed to an oocyte with more than 100 spermatozoa attached to oocyte ZP, Level 3 if the oocyte has between 50 and 100 bound spermatozoa to its ZP, Level 2 for 20 to 50 bound spermatozoa and Level 1 if the oocyte has less than 20 bound spermatozoa (Figure 6A). Figures 6B and 6C show the temporal evolution of the rate of oocytes associated to each level for the group of penetrated unfertilized oocytes, and the group of fertilized oocytes respectively. For the unfertilized oocytes, we observe that the proportion of oocytes in each level changes little over time despite multi-penetration of the oocytes showing that penetration has no significant effect on the adhesion properties of the ZP. The story is different when fertilization occurs. Indeed, Figure 6C shows that the oocytes from the group of fertilized oocytes maintain their initial level of bound spermatozoa until fertilization take place then start to switch to lower and lower levels (i.e. less and less bound sperm) during typically 1 hour. As for ZP permeability decrease, gamete fusion (and not penetration) appears as the trigger of sperm de-adhesion from the ZP. Despite the poor accuracy in the quantification of the ZP de-adhesion kinetics compared to the ZP permeability inhibition kinetics, we can reasonably say that both permeability and adhesion inhibitions of the ZP occur within the same timeframe. This raises the question of whether loss of adhesion and loss of permeability are two independent manifestations of a fertilization-triggered process, or whether loss of penetrability is directly the consequence of the decrease in ZP-bound spermatozoa due to de-adhesion. The fact that the group of penetrated but unfertilized oocytes shows no correlation between the number of ZP-bound spermatozoa and the occurrence of penetrations argues in favor of two distinct manifestations.

**Figure 6:**
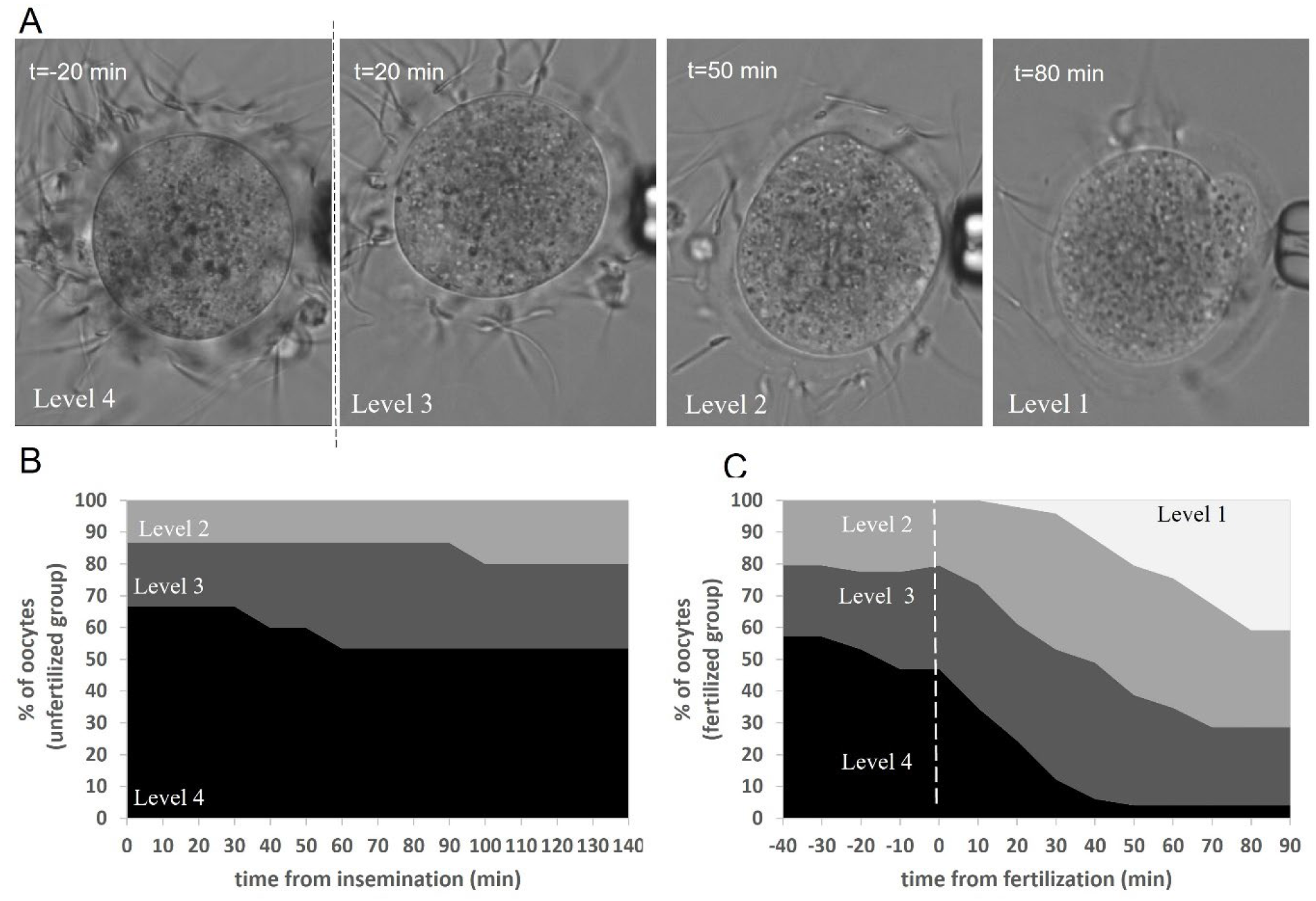
Evolution of the number of spermatozoa bound to the ZP ff an oocyte with its penetration and fertilization status. A-Typical sequence of an inseminated oocyte 20 min before fertilization, 20 min, 50 mn and 80 min after fertilization. Levels 4, 3, 2 or 1 respectively correspond to oocytes with (i) more than 100 spermatozoa, (ii) between 50 and 100 spermatozoa, (iii) between 20 and 50 spermatozoa, (iv) less than 20 spermatozoa bound to the ZP. B-Temporal evolution of the proportion of oocytes in each level for the group of penetrated but unfertilized oocytes. C-Temporal evolution of the percentage of oocytes associated to each level for the group of fertilized oocytes. The dashed white line indicates the time of fertilization.

### The choreography of a penetrating spermatozoon reveals its capacity to fertilize or not

Fertilization success/failure necessarily results from favorable/unfavorable interactions between the penetrating spermatozoon and the OPM. We took advantage of the rounds of observation of the oocytes in kinetic *in vitro* fertilization experiments to describe how fertilizing and non-fertilizing spermatozoa interact with the OPM once penetrated, in order to characterize the role of the different components of the oocyte (i.e. ZP, PVS, OPM) and of the spermatozoon (i.e. head, flagellum) in promoting fertilization and preventing polyspermy.

#### Typical 3-phase choreography of successful fertilizing spermatozoa

When crossing the ZP and entering the PVS of an oocyte, the penetrating spermatozoon is generally oriented perpendicular to the ZP and thus to the OPM (Video 1) or slightly tilted (Video 2). Its flagellum beatings allow the spermatozoon to make its way through the ZP with its head gradually entering the PVS (Video 1, penetration flag). Because the width of the PVS at the spermatozoon emerging site is often of the order the spermatozoon head length, its tip rapidly bumps into the OPM delaying the progression of the spermatozoon through the ZP.

The spermatozoon may stagnate in this position perpendicular to the oolemma for several minutes (∼6 min in Video 1, wiping flag) with its head oscillating in a wiping motion, its tip grazing the surface of the oolemma (Figure 7A). When, by dint of oscillations, the spermatozoon manages to disengage a bit more from the ZP, its head can tilt more and more parallel to the OPM and starts to adhere to it. The bonds that hold the spermatozoon head to the OPM transform its wiping motion into a rolling motion (Video 1, crawling/rolling flag). These bonds are obviously sufficient to prevent the spermatozoon head from moving away from the OPM despite flagellum movement, but still sufficiently labile so that the sperm head can slowly crawl on the OPM as its flagellum continues to progress through the ZP) (Figure 7A). This crawling/rolling regime can last for several minutes (∼2,5 min in Video 1 crawling/rolling flag) before the adhesion of the equatorial segment of the spermatozoon with the OPM becomes strong enough to prevent it from crawling further despite the persistent thrust force of its flagellum oscillations (Video 1 & Video 3, rolling/pushup flag). In that case, both the adhesion that maintains the spermatozoon head laying on the OPM and the ZP that maintains its flagellum trapped, impose limit constraints to flagellum oscillations in the PVS and thus to the spermatozoon head whose rolling may change into a mix of rolling and pushup-like movements on a stable area of the OPM (Figure 7A). This regime generally lasts for one to few minutes (∼1 min in video 1, rolling/pushup flag) and ends in the fusion of the gametes, identified by the sudden stillness and stiffness of sperm flagellum (Video1 & Video 2, white star fertilization flag).

**Figure 7:**
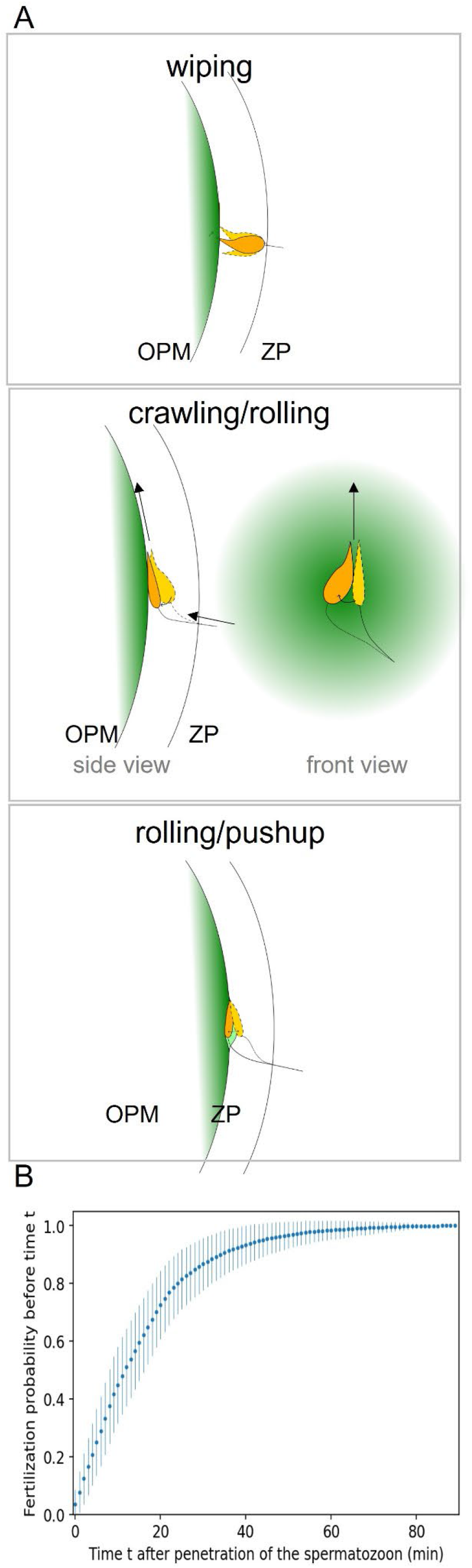
From penetration to fusion: phases and kinetics. A-3 interaction phases. Wiping phase: the spermatozoon head bumps into the OPM, its head oscillates in a wiping motion, its tip grazing the OPM. Crawling/rolling phase: the spermatozoon progresses in the PVS crawling on the OPM, its head having a rolling motion. The arrows indicate the progression of the flagellum through the ZP and the progression of the head on the OPM. Rolling/pushup phase: the adhesion of the spermatozoon head to the OPM is strong enough to stop its crawling on the OPM and the progression of the flagellum though the ZP. The oscillation of the flagellum limited by the adhesion of the sperm head on one site and the entrapment of the flagellum on the other side, imposes to the head the typical rolling/pushup motion leading to fusion. B-Fertilization after penetration time cumulative distribution of the fertilizing spermatozoon in fertilized monospermic oocytes.

Alternatively, it happens that during the crawling/rolling phase, the sperm head progresses far enough from its ZP entry site for it to fully enter the PVS (Video 4 & Video 5 red star full penetration of flagellum flag). Once released from the ZP, the flagellum can oscillate more freely, limited only by the walls of the PVS (i.e. the OPM and the ZP), which do not present strong limitations compared to its entrapment in the ZP. When fully penetrated in the PVS, we noticed that the only spermatozoa able to fertilize are those with medium amplitude flagellum oscillations, as in Video 6, allowing their heads to adopt a stable position on the OPM, and displaying the rolling/pushup oscillations preceding fusion, similar to the fertilizing spermatozoa trapped in the ZP.

This direct observation of the progression of fertilizing spermatozoa in the oocyte, from penetration to fusion therefore reveals a typical journey ranging from a few minutes to often more than 15 minutes, involving up to 3 phases of gamete interaction (i.e. wiping, crawling/rolling and rolling/pushup), the last of which is decisive for fusion. From the observed penetration and fertilization time windows of the first fertilizing spermatozoa of the 54 monitored monospermic oocytes (Figure 3A) and applying a statistical treatment of these events relying on a uniform probability assumption in each time window, we inferred the cumulative distribution function of the fertilization event vs time after penetration of a first fertilizing spermatozoon (Figure 7B). The average fertilization time of a first fertilizing spermatozoon after its penetration in the PVS of an oocyte can be deduced: 15.8 ± 5.7 minutes.

#### Unfavorable flagellum oscillations of a penetrated spermatozoon prevent it from fertilizing

Coupling acrosome reacted spermatozoa with oocytes freed from their ZP, we previously showed that a specific flagellum beating that produced a push-up like movement of the spermatozoon head on the OPM was mandatory for fusion to occur (Ravaux et al., 2018, 2016). The present results show that this movement is also an essential parameter for fusion in physiological ZP intact conditions. Remarkably, this pro-fusion oscillation is observed for any spermatozoon that penetrates a not yet fertilized oocyte as long as its movement is constrained by the adhesion of the sperm head with the OPM on one side, and by the entrapment of its flagellum in the ZP on the other side. When the latter condition is not met anymore, which happens when the flagellum completely penetrate the PVS (Video 4, Video 5), this favorable oscillation sometimes transforms into an erratic movement as illustrated in Video 4, causing the spermatozoon head to rotate on the OPM and therefore preventing the stability necessary for fusion. In such cases, although the oocyte is not yet fertilized, the spermatozoon does not manage to fuse including when it is the first penetrated spermatozoon. Instead, it typically detaches from the OPM after a variable period of time and does not reattach or only transiently, confirming that inappropriate flagellum movements lead to fertilization failure, and explaining why a first penetrated spermatozoon may not fertilize.

#### The ZP promotes fusion by channeling flagellar movements

We observed that as long as a penetrating spermatozoon has its flagellum trapped in the ZP and its head adhering to the OPM, it cannot adopt movements unfavorable to fusion, since these require freedom that is not allowed geometrically due to the constraints imposed by the trapping of its flagellum in one side and the adhesion of its head to the OPM on the other side. Conversely, from the moment the whole flagellum enters the PVS, the ZP constraint disappears and, the flagellum becomes free to adopt a more erratic beating pattern that is not conducive to fusion. This reveals that the ZP plays a key, hitherto unsuspected role in mechanically constraining the spermatozoon to adopt movements conducive to fusion. We examined the position of all penetrated fertilizing and unfertilizing spermatozoa over the rounds of observation of the oocytes. Of the fertilizing spermatozoa (60 fertilizing spermatozoa), a large majority (73%) fuse while their flagella are still trapped in the ZP, probably reflecting for many of them the favorable role played by the trapping of its flagellum in the ZP. Conversely, all non-fertilizing spermatozoa end up fully penetrated in the PVS. This does not mean that all these spermatozoa fail to fertilize due to an inadequate beating (especially those that penetrated after fertilization that have other reasons for failing to fertilize, as we will see later) but we do know that some of them adopt an erratic beating dooming them to fertilization failure.

#### Impaired interaction between penetrating spermatozoa and the plasma membrane of fertilized oocytes

By contrast to the almost systematic adhesion observed when a spermatozoon penetrates an unfertilized oocyte, we often observed that the spermatozoa penetrating a fertilized oocyte do not firmly attach to the OPM. As illustrated in video 7 corresponding to a fertilized oocyte, the adhesion of the penetrating spermatozoon during the crawling/rolling phase is weak enough to not be a hindrance to its progression through the PVS. As soon as its flagellum is released from the ZP, the spermatozoon starts to swim in the PVS around the oolemma without adhering to the OPM anymore. Consistently, when observed during the rounds of observation of their oocytes, most of the unfertilizing spermatozoa present in the PVS of fertilized oocytes are swimming around the OPM and few are attached to the OPM. A small adhesion to the OPM therefore appears to be the main cause of the fertilization failure of the spermatozoa penetrating the PVS of an already fertilized oocyte. This direct observation is consistent with the numerous studies of fertilized oocytes, reporting spermatozoa swimming in the PVS without being able to adhere to and fuse with the OPM. This lack of adhesion is generally speculated as being due to the loss of the OPM affinity for the spermatozoa. However, the possibility of the reverse -that this impairment in gamete adhesion is due to an inhibition of the affinity of the spermatozoon for the OPM while penetrating the PVS of fertilized oocyte-cannot be ruled out objectively and requires further investigation.

#### Probable neutralization in the PVS of the spermatozoa penetrating a fertilized oocyte

CD9 and JUNO are the two oocyte proteins identified to date as essential for fusion in mice (Bianchi et al., 2014; Kaji et al., 2000; Le Naour et al., 2000; Miyado et al., 2000). We previously showed that 50% of the oocyte membrane CD9 is suddenly released from the OPM four minutes after fertilization (Ravaux et al., 2018) and although the precise kinetics of release of JUNO are yet unknown, oocytes are shown to be significantly depleted from the fertilized OPM within one hour after fertilization (Bianchi et al., 2014). This post-fertilization shedding process was proposed to impair both the affinity of the OPM for spermatozoa through the depletion of these proteins and the affinity of the penetrating spermatozoa to OPM through the binding of JUNO released in the PVS to its sperm receptor IZUMO1 (Bianchi et al., 2014). However, this hypothesis has not been studied further since then. Here, standard *in vitro* fertilization assays performed with CD9-EGFP oocytes (N=80) reveals that a significant amount of CD9 and very little JUNO remain at the OPM of fertilized oocytes which is consistent with previous studies (Bianchi et al., 2014; Ravaux et al., 2018). Additionally, we noticed that JUNO and CD9 are still massively present in the PVS of fertilized oocytes 4 hours after insemination of the oocytes while the both proteins remained located at the OPM of the control unfertilized oocytes (Figure 8A). This shows that once released from the OPM, CD9 and JUNO cannot easily cross the ZP and therefore accumulate in the PVS. The consequence is that a spermatozoon entering the PVS after a first fertilization must evolve in a CD9 and JUNO rich-environment, and this released content may interact with it. We have investigated this possibility, and interestingly observed that the heads of all penetrated unfertilizing spermatozoa were systematically covered with JUNO and CD9 (Figure 8B). This strongly supports the idea that the post-fertilization release of CD9 and JUNO can inhibit the fusing ability of the OPM because of the depletion of JUNO and CD9, and neutralize the new penetrating and already penetrated spermatozoa by binding to their head, inhibiting their capacities to properly interact with the OPM and fuse. In this case, the ZP would take on an additional role that is essential for blocking polyspermy: that of an exit barrier that prevents the EPV components that neutralize the spermatozoa in the PVS from diluting outside.

**Figure 8:**
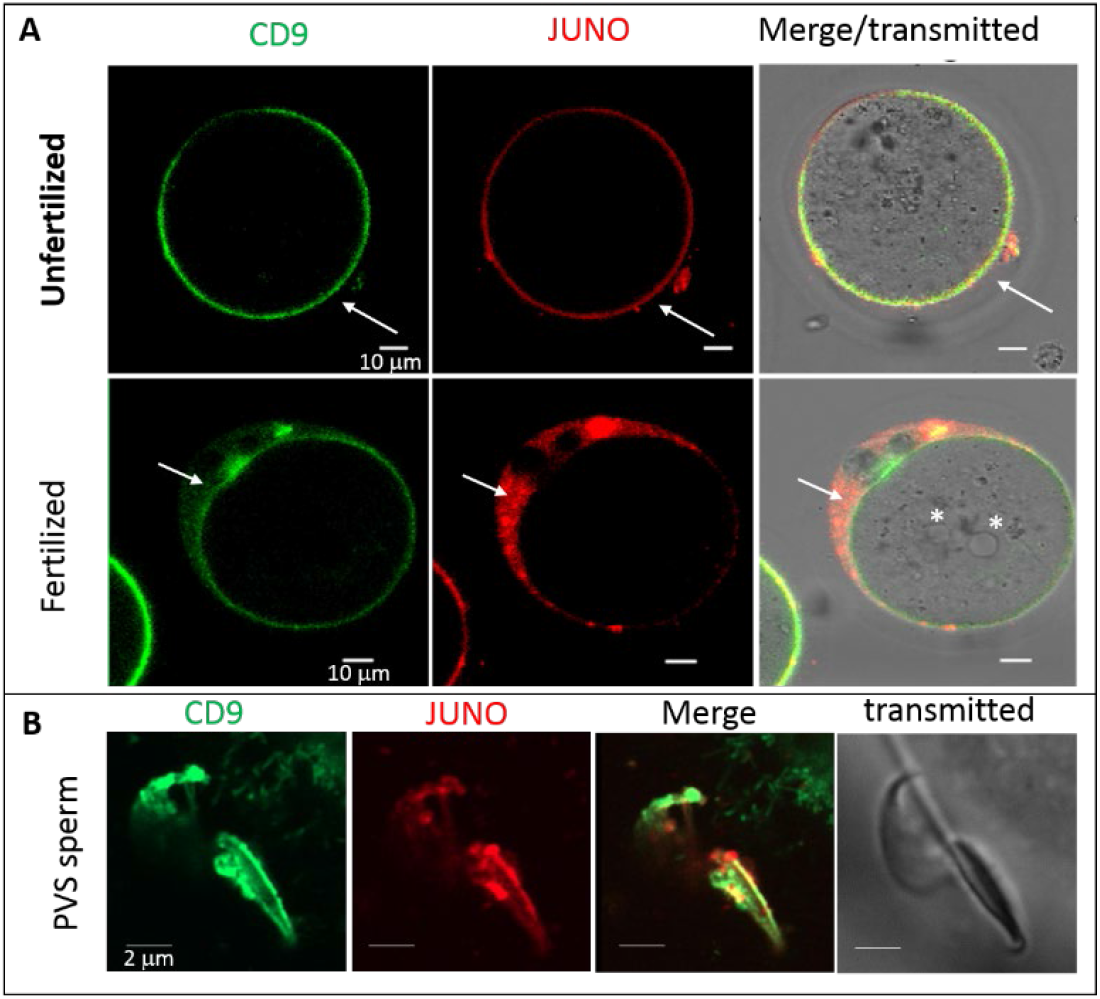
Persistence in the PVS of JUNO and CD9 released after fertilization and presence of CD9 and JUNO on the heads of the sperm in the PVS. A-Confocal images of unfertilized and fertilized CD9-EGFP (green) oocytes stained with Juno-dylight 633 antibody (red) 4 hours after insemination. The white arrows designate the position of the PVS. The white stars show the 2 pronuclei resulting from monospermic fertilization. B-Confocal images of spermatozoa penetrated in the PVS of CD9-EGFP oocytes stained with Juno-dylight 633 antibody, 4 hours after insemination.

## Discussion

### Secrets of successful fertilization: pushup oscillations with or without the help of the ZP

From penetration to fertilization, the interaction between the penetrating spermatozoon head and the OPM generally takes place in 3 phases: 2 approach phases and 1 actual fertilization phase (Figure 7). The first approach phase enables the sperm to lay its head on the OPM. In the second phase, the spermatozoon slowly crawls over the OPM until it develops an adhesion strong enough to prevent it from going any further. For fertilization to occur, the spermatozoon head must adopt in the last phase, an oscillating push-up like movement on the OPM, whether or not its flagellum is still trapped in the ZP. Each phase can last from less than a minute to several minutes, and the whole process can exceed 15 minutes. It is worth noting that the last phase was shown to fail in all cases where spermatozoa entered the PVS more than 15 minutes after fertilization due to impaired adhesion. It is worth noting that it also failed in many cases where spermatozoa entered the PVS before or shortly after another spermatozoon fertilized, but could not fertilize before completion of the fertilization block. A long time spent in the PVS without fertilizing around the time of a first fertilization can therefore cause its fertilization failure.

This study shows that the first penetrating spermatozoon is not necessarily the one that fertilizes. The fact that it contradicts an almost century old dogma is not the most important point of this discovery, especially as this dogma still reflects the majority of the fertilization cases. More interesting is the reason why a first penetrating spermatozoon, often presented as “the best”, can fail to fertilize. The sole difference observed between first penetrated fertilizing and non-fertilizing spermatozoa is the motion pattern of their flagella. As long as a piece of their flagellum remains trapped in the ZP, no distinctions can be done from one spermatozoon to another regarding its interaction with the OPM: the trapping of the flagellum in the ZP in one side, and the adhesion of the head to the OPM in the other constrain the penetrating spermatozoon to adopt the motion pattern compatible with successful fertilization. This evidence extends to the physiological context of ZP intact oocytes, the necessity to have a specific flagellum beating mode for fusion previously that was demonstrated with ZP-free oocytes (Ravaux et al., 2018, 2016). But here, the new and exciting discovery is the crucial contribution of the ZP to promote fusion by channeling flagellar movements.

### Block to penetration: ZP block is too slow to significantly contributes to prevent polyspermy but the low natural ZP permeability does

We determined for the first time the natural permeability of the ZP to spermatozoa in mice and found that it is low, allowing only one spermatozoon to pass through every 47 minutes on average. After the first fertilization, the ZP undergoes a progressive and persistent loss of its permeability with a time constant equal to 48 min. This is in good agreement with literature (Baibakov et al., 2007; Barros and Yanagimachi, 1971; Bleil et al., 1981; Burkart et al., 2012; Fahrenkamp et al., 2020; Miller et al., 1993; Nishio et al., 2024; Que et al., 2017; Tokuhiro and Dean, 2018). As the ZP block prevents spermatozoa from penetrating and thus from fertilizing oocytes, it is generally associated to the term polyspermy which has led to the expression “ZP block to polyspermy”. However, as shown in Figure 9, at the post-fertilization time *t*_*PF*_ = 6,2 ± 1.3 minutes corresponding to the time constant of the polyspermy block, the difference between the mean number of spermatozoa per oocyte actually penetrated after a first fertilization and what it would have been in the absence of any fertilization-induced decrease in ZP permeability (i.e. without ZP block) is still negligible. This shows that during the post-fertilization period when a penetrating spermatozoon can still fertilize, fertilization-induced ZP block does not add value yet to the initial low natural ZP permeability. It will do after, and there is no doubt that the ZP block eventually makes the ZP permanently impenetrable to spermatozoa, but the slowness of the process makes it useless for preventing polyspermy. This is in agreement with experiments showing that in mouse oocytes missing ovastacin (i.e. the protease responsible for the cleavage of ZP glycoprotein ZP2 resulting in the ZP block), oocytes remain monospermic despite the fact that ZP-block cannot take place (Nozawa et al., 2018). The often-used expression “ZP block to polyspermy” is therefore inappropriate and misleading for mice. Conversely, the low natural ZP permeability to spermatozoa makes it a key asset in preventing polyspermy, by limiting the number of penetrations during the entire period (before and after the first fertilization) when oocytes are fertilizable and the risks of polyspermy are real.

**Figure 9:**
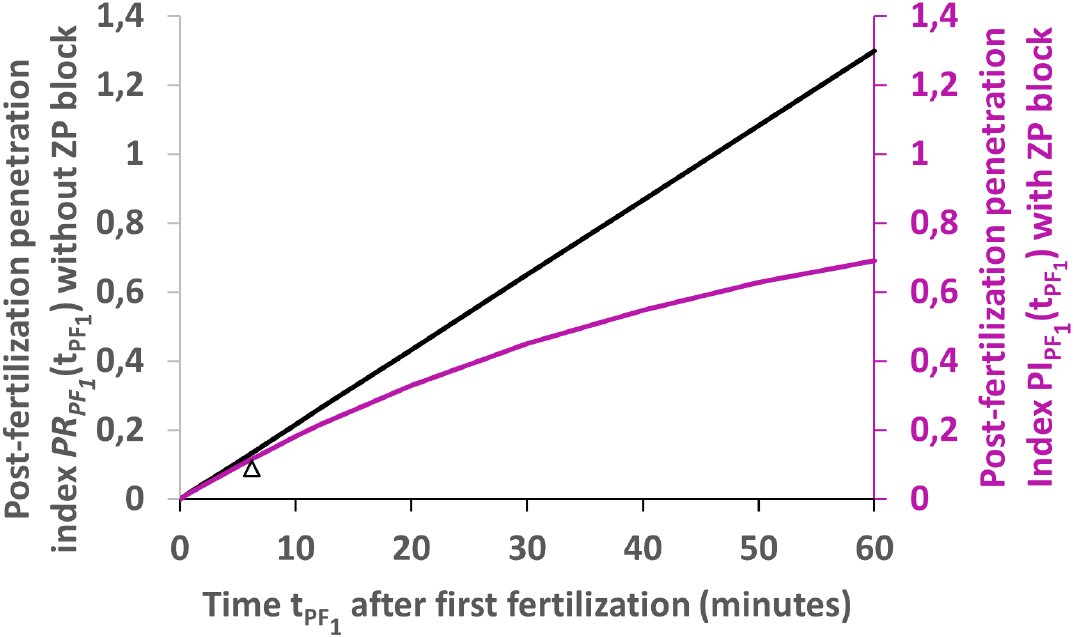
Evolution of post-first fertilization penetration index, in the hypothetical case of the presence of ZP block (purple) and in the hypothetical case of the absence of ZP block (black). The arrow head indicates the absence of significant difference between both situations at time *t*_*PF*_ = 6,2 min corresponding to the time constant of the fertilization block.

### Block to polyspermy: PVS block should be the missing fast contribution to polyspermy prevention

Our observations reveal fast kinetics of the polyspermy block, with time constant equal to 6,2 ± 1.3 minutes, triggered by the first fertilization, and preventing new penetrating and already penetrated spermatozoa to fertilize. The fertilization failure of spermatozoa penetrating the PVS of an already fertilized oocyte is generally justified by the membrane block, resulting in a progressive and persistent loss of OPM ability to fuse with additional spermatozoa, taking place along with an inhibition of the OPM ability to bind spermatozoa (Gardner et al., 2007; Kryzak et al., 2013; Maluchnik and Borsuk, 1994; Wolf, 1978) that have been observed on fertilized ZP-free oocytes. The molecular origin of this membrane block, has not yet been elucidated, but the post-fertilization time after which the OPM of ZP-free oocytes is no longer able to fuse with spermatozoa, was assessed typically to one hour. Although such a membrane block is consistent with our observation of the low adhesion between the OPM of fertilized oocytes and a penetrating spermatozoon (Video 7), the time after which it is no longer able to fuse with spermatozoa, assessed typically to one hour, is not compatible with the fast polyspermy block of a few minutes that we have determined. A major difference between previous studies on the membrane block and our study is the physiological presence of ZP and therefore PVS in our case, suggesting that the missing rapid contribution to the polyspermy block could come from the PVS. Our study shows that the recently highlighted post-fertilization release of CD9 and JUNO (Bianchi et al., 2014; Ravaux et al., 2018) can potentially create a molecular framework for both membrane and PVS blocks, the former by depleting the OPM of two proteins mandatory for gamete fusion success, the latter by loading the PVS with these molecules. We have demonstrated that CD9 and JUNO accumulate in the PVS after the first fertilization and are still there 4 hours after fertilization (Figure 8A). Moreover, we have shown that the heads of unfused spermatozoa in the PVS are systematically coated with these released proteins (Figure 8B). Our hypothesis for the PVS-block, consistent with the observed low adhesion between the OPM of fertilized oocytes and a penetrating spermatozoon, is that JUNO and CD9 passivate the spermatozoon by inducing non-specific steric hindrance of the heads of the penetrated spermatozoa and/or specific inhibition of sperm ligands like IZUMO1. Each of these possibilities can contribute in neutralizing spermatozoon ability to properly adhere and fuse with the OPM. With regard to the respective kinetics of membrane*-* and PVS*-* blocks, although both would rely on a common process of release of OPM molecules after first fertilization, we do not expect them to be fully effective in the same time scale. We would expect from the kinetics of loss of OPM receptivity to spermatozoa-established by experiments on ZP-free oocytes-(Kryzak et al., 2013; Maluchnik and Borsuk, 1994) that the membrane*-*block requires the OPM to be almost completely released from JUNO (∼1 hour (Bianchi et al., 2014)) to become fully effective. In contrast, we expect that the *PVS-*block requires only partial release of CD9 and JUNO, achievable in a few minutes, for complete passivation of the penetrating spermatozoa. The reasoning behind this assumption is that CD9 and JUNO are released, diffuse, accumulate and remain in the tiny confined volume of the PVS (∼35.10^−6^ μL) (Figure 8A), and that a small fraction of what is ultimately released is sufficient to rapidly coat the head of the spermatozoon penetrating in the PVS or already there, since spermatozoon head surface area is typically 1/1000 of the surface area of the OPM. This suggests that passivation of the penetrated spermatozoa can rapidly occur after the onset of the release process triggered by fertilization, and therefore that the *PVS-* block, relying on neutralization of the spermatozoa in the PVS, may be the fast polyspermy block we inferred. It is worth noting that the ZP would play an indirect but crucial role in this process as an impenetrable exit barrier for the components released by the OPM after first fertilization, trapping them in the PVS where they accumulate, creating an unfavorable environment for new fertilizations.

## Conclusion

By questioning notions that have been dogma for decades, such as the role of the ZP in fertilization and block to polyspermy, or the omnipotence of the first spermatozoon to penetrate the oocyte, this kinetic study forces us to move away from classical views. Our observations provide new pieces of evidence that gamete interaction and fusion do not depend only on protein factors and that mechanical factors involving the ZP and influencing gamete membrane interaction are also crucial. Which mechanical parameters come into play and how they contribute to locally create the physico-chemical environment prone to fusion are multidisciplinary questions that require the development of interfacial synergetic research to progress in the understanding of the gamete fusion process. Our results also show that the penetration block is first controlled by the low permeability of the ZP and then by the fertilization triggered ZP-block. However, because of the slowness of the ZP-block, only the initial low permeability of the ZP (but not the ZP-block) can contribute to prevent polyspermy. If ZP-block cannot contribute in blocking polyspermy, a relevant research path is to investigate its actual function in the fertilization and/or development process. As for the reason why penetrated spermatozoa fail to fuse after a first fertilization, it becomes clear that it cannot be based solely on the too slow membrane block. Other faster contributions are required to account for the fastness of the polyspermy block. The discovery that the penetrated non-fertilizing spermatozoa are coated with released oocyte suggests that the missing piece of the block to polyspermy puzzle consists in a fast neutralization of any new penetrating and already penetrated spermatozoa in a PVS that has become a sperm inhibiting medium. The missing fast block would therefore be a PVS block. This model opens a new avenue on a field of research that have been at a standstill for a couple of decades: how polyspermy is prevented in mammals? At a time when infertility issues are becoming a real concern, understanding the still unresolved questions about how fertilization works will permit to address these infertility issues with the greatest discernment possible.

## Material and Methods

### In vivo fertilization

8 to 12-week-old wild-type (WT) female mice (B6cbaF1 background) were super-ovulated by intraperitoneal injections, first of 5 IU PMSG, followed by 5 IU hCG 48 h apart and immediately mated with a male. 15 hours after mating, plugged females were sacrificed, oocytes were recovered, washed, fixed and stained with Hoechst and imaged in confocal microscopy to determine the number of penetrations and fertilizations of each oocyte.

### Sperm preparation for in vitro fertilization experiment

Sperm was obtained from WT male mice (B6cbaF1 background). Sperm were expelled from cauda epididymis and vas deferens into Ferticult® IVF medium (Fertipro, France) under mineral oil. Sperm were then incubated in Ferticult® at 37°C, 5%CO2 in air for 1.5 h to induce capacitation.

### Oocytes preparation for in vitro fertilization experiment

*CD9-EGFP* and *Wild-Type* oocytes were obtained from 6 to 12-week-old female mice (B6cbaF1 background for WT mice and C57black6J background for CD9-EGFP transgenic mice oocytes (Miyado et al., 2008)). Female mice were super-ovulated as previously described. Cumulus/oocytes masses were collected into a Ferticult® IVF medium drop 14 h later by tearing the oviduct ampulla from sacrificed mice.

### In vitro kinetics and standard experiments

Both kinetics and standard *in vitro* experiments started by insemination of cumulus/oocytes masses recovered from super-ovulated females, with capacitated spermatozoa at the concentration of 10^6^ spermatozoa/mL and put in an incubator (37°C, 5% CO2) .15 minutes of such co-incubation were generally sufficient for most of cumulus cells to detach and for most oocytes to be surrounded by numerous spermatozoa bound to their ZP without being penetrated or fertilized yet. For kinetics *in vitro* experiment, part of the oocytes was isolated with associated bound spermatozoa in individual drops (to be able to identify each of them) 15 minutes after insemination, successively immobilized by gentle pipette contention and filmed in close-up in turn during typically 4 hours. Standard *in vitro* experiment consisted in maintaining the remaining oocytes in co-incubation with of 10^6^ spermatozoa/mL in the incubator for 4 hours. Both kinetics and standard experiments were stopped 4 hours after time of insemination, the oocytes were washed from the spermatozoa still attached to their ZP, keeping the identity of the oocytes previously studied in kinetics experiments, and imaged in confocal microscopy after DNA staining. 10 independent *in vivo* experiments were performed for a total of 211 oocytes. 11 independent *in vitro* experiments were performed for a total of 93 isolated (kinetics experiments) and 220 control oocytes.

### Determination of the number of penetrations and fertilizations

4 hours post-insemination, all the oocytes (from standard and kinetics *in vitro* experiments) were washed to free them from the spermatozoa still attached to the ZP, fixed, stained with Hoechst, and imaged with a confocal microscope. The number of spermatozoa that successfully crossed the ZP and the number of penetrated spermatozoa that fertilized were determined for each oocyte on the basis on the number of DNA spots of sperm nuclei in the PVS (penetrated non fertilized eggs) and on the number of decondensed sperm nuclei and/or male pronuclei in the oocyte cytoplasm (penetrated fertilized sperm) respectively.

### Determination of the penetrating and fertilizing status of spermatozoa

The few oocytes isolated 15 min post-insemination with the spermatozoa attached to their ZP from the insemination chamber were imaged in bright field in turns at 37°C, by successively immobilizing them at the tip of a micropipette through gentle aspiration. When immobilized, the whole volume of the oocyte was explored by tuning vertically the objective (and therefore the focus plane) in order to screen plane by plane the oocyte and associated spermatozoa, looking for penetrating and fusing sperm. Triple checking was done by changing twice the position of the oocyte at the tip of the pipette in order to obtained 3 different angle views of the oocyte. The numbers of penetrated and fused spermatozoa were counted at each round of observation of spermatozoa and compared to the previous round, allowing to determine the order of penetration of each spermatozoon into the PVS of a given oocyte and its fertilization status. A spermatozoon was considered penetrated when at least its head was emerged from ZP into the PVS. A spermatozoon was further considered fused when its flagellum was immobile and straight, as pinned in the OPM and its head lying on the OPM or partially or totally engulfed in the oocyte.

### Determination of the penetration times windows (Figure 3A & 3B)

When the penetration of a spermatozoon occurred during a round of observation of the oocyte, the post-insemination time at which the spermatozoon penetrated was obtained directly to the minute. By contrast when the spermatozoon penetrated between two rounds of observation of the associated oocyte (i.e between rounds N-1 and N for instance), we could just deduce that penetration occurred between the end of round N-1 and the beginning of round N without more precision. In that case, we obtain a penetration time window symbolized by a line of appropriate length in Figure 3. All times of the time window were considered as equiprobable possible penetration times.

### Determination of the fertilization time windows (Figure 3A)

Only a spermatozoon whose flagellum is still beating and whose head equatorial segment is interacting with the OPM can fertilize. We have previously shown that the onset of gamete fusion and the permanent stop of the flagellum beating of the involved spermatozoon are concomitant to within one minute [Ravaux 2016]. Therefore, when by chance the flagellum arrest occurred during an observation cycle, this event was considered a precise marker of the time of fusion. On the other hand, when oocyte fertilization occurred between two rounds of oocyte observation (between rounds N-1 and N, for example), the precision of the fusion time was less. However, unlike the evaluation of the penetration time, which could only be as precise as the time interval between the end of round N-1 and the start of round N, the determination of the fusion time could be refined on the basis of the studied kinetics of sperm engulfment by the fertilized oocyte and then that of PB2 release (Figure 2). This was done by determining for each round of observation of an oocyte whether or not it was fertilized at post-insemination time of observation *t*_*PI*_, and if so, by assessing its post-fertilization stage at this time, from which an estimate of fusion time *t*_*F*_ was derived. 3 stages of fertilization were considered:

*Stage 1-sperm engulfment* (the head of the fused sperm is not visible at the OPM 24±3 min after fusion)

-If the head of the fused sperm was still visible on the OPM, then fusion was considered to have taken place at: *t* _*PI*_ − 27 *min* < *t*_*F*_ < *t*_*PI*_

-If the head of the fused sperm was not seen anymore, fusion was considered to have taken place at: *t*_*F*_ < *t*_*PI*_ − 21 *min*

*Stage 2-PB2 release Phase 1* (angle between the OPM and PB2 higher than 90°, beginning: 28± 2min after fusion; end: 49± 6min after fusion)

-If the PB2 was observed in Phase 1, fusion was considered to have taken place at: *t*_*PI*_ − 55 *min* < *t*_*F*_ < *t*_*PI*_ − 26 *min*

Stage 3 PB2 release Phase 2 (angle between the OPM and PB2 smaller than 90°; beginning: 49± 6min min after fusion; end: 73± 10 min after fusion)

-If the PB2 was observed in Phase 2, fusion was considered to have taken place at: *t*_*obs*_ − 83 *min* < *t*_*F*_ < *t*_*PI*_ − 43 *min*

-If the PB2 was observed completed at *t*_*obs*_, fusion was considered to have necessary taken place at : *t*_*F*_ < *t*_*PI*_ − 63 *min*

As an oocyte may have been observed before it was fertilized but also during each of these 3 fertilization stages, we could have up to 4 determinations of the fusion time intervals. Unless a stage was misidentified, all the intervals determined were expected to have a common overlap (which turned out to be the case) corresponding to the more accurate estimate of the fusion time window we could obtain. Within this fusion time window, all times were considered as equiprobable possible fusion times. They are symbolized by the black segments of appropriate length in Figure 3A

#### Statistical treatment of penetrations and fertilizations chronograms to study penetration and fertilization blocks

Our experiments enable to determine time windows for the penetration and fertilization events of each spermatozoon. To deduce averaged fertilization, penetration and polyspermy indexes and their confidence intervals shown on figure 5, we have implemented a statistical treatment of these data based on the simple assumption that each event takes place at a given time in its observed time window with a uniform probability distribution. Based on this hypothesis, we have numerically computed probability distributions of numbers of spermatozoa penetrated or fertilized for each oocyte on timescales whose zeros are shifted either to the first penetration (Figure 5B) or first fertilization (Figure 5A, C) event. From these distributions, averaged penetration, fertilization or polyspermy indexes are deduced. Python scripts used for these computations can be provided on demand by the corresponding author.

#### Localization of CD9 and JUNO by confocal imaging of CD9-EGFP oocytes and penetrated spermaotozoa

4 hours after insemination of cumulus/CD9-EGFP oocyte masses with spermaotozoa (10^6^ spermatozoa/mL), the oocytes were washed to free them from bound spermatozoa. The oocytes were loaded with Hoechst for DNA staining and stained with anti-JUNO/FOLR4 monoclonal antibody (BioLegend) coupled with a Dylight 633. Samples were imaged with Leica SP5 confocal microscope or Zeiss Airy scan microscope on the lookout of the localization of CD9 and JUNO at the OPM, in the PVS and on the prenetrated spermatozoa.

## Supplementary information

The Videos can be found online at: https://idata.phys.ens.fr/index.php/s/smxCQfzEQoeqYpe

## Ethics Statement

All animal experiments were performed in accordance with national guidelines for the care and use of laboratory animals. Authorizations were obtained from local (Animal Care and Use Committee Charles Darwin, France (#30204)) and governmental ethical review committees via APAFiS Application (Autorisation de projet utilisant des animaux à des fins scientifiques), authorization APAFIS #30204-2021030415199914 v3.

## Author contributions

Conceived and designed the experiments: YD, SF, CG. Conceived the statistical analysis: NR. Performed the experiments: YD, SF and MO. Analyzed the data: YD, NMF, SF, NR, CG Contributed reagents/materials/analysis tools: YD, SF, NMF, DS, MO, EP, SB, AZ, NR, CG. Wrote the paper: NR, CG.

## Acknowledgments

The authors warmly thank Eleonore Touzalin and Gwendoline Firmin for animal care, Frédéric Pincet and Sophie Cribier for fruitful discussions.

